# Proteomic profiling of drug and nutrient transporter expression by small intestinal region in neonates, pediatrics, and adults

**DOI:** 10.64898/2026.06.13.731200

**Authors:** Vicky L. Sun, Kai Wang, Edward J. Kelly, Samuel L.M. Arnold

## Abstract

Transporters localized to the small intestinal epithelium govern the absorption of many orally administered drugs and nutrients, and thorough characterization of transporter protein abundance is key to accurate prediction of small molecule absorption. Previous studies have reported more than 100 solute carrier and ATP-binding cassette superfamilies expressed in the intestine, but only a small subset of these transporter proteins have been quantified in native intestinal tissue. In addition, a majority of published transporter abundance data for the intestine was collected with adult tissue. Here, we report the identification and relative quantification of 87 solute carrier and 18 ATP-binding cassette superfamily transporters by intestinal region in neonatal, pediatric, and adult small intestinal mucosa using data-independent acquisition mass spectrometry. This study identified transporters uniquely expressed by intestinal region in all three age groups and region-associated abundance for 23 SLC/ABC transporters in pediatrics and adults. The protein abundance data reported here provide a valuable resource for the prediction of nutrient and drug absorption and efflux across age and will inform future studies of specific transporters and age-associated differences in transporter abundance.

**Significance statement:** We report the relative abundance of key xenobiotic transporters along the length of the small intestine in neonates, pediatrics, and adults based on data-independent acquisition mass spectrometry. Intestinal region-associated transporter expression was observed for drug transporters and under-studied nutrient transporters across age groups.

## Introduction

A primary function of the small intestinal epithelium is to facilitate the absorption of nutrients from the diet while serving as a protective barrier against the entry of potentially toxic xenobiotics.^1,2,3,4,5^ The large surface area of the small intestine contributes to nutrient and xenobiotic absorption: circular folds (plicae circulares) in the duodenum and jejunum augment absorptive area while microscopic protrusions known as villi and microvilli line the epithelium throughout the small intestine.^1,6,7,8^ The intestinal epithelium is a monolayer of various cell types including enterocytes, which absorb many small compounds by paracellular or transcellular diffusion.^1,2,9^ Due to the presence of tight junctions, many hydrophilic and/or large drugs and nutrients depend on transporter proteins localized to the plasma membrane of enterocytes to assist their passage to systemic circulation.^3,10,11,12,13,14^

Many transporters critical for nutrient and drug absorption belong to the solute carrier (SLC) or ATP-binding cassette (ABC) superfamilies. To date, 395 SLC and 48 ABC transporter genes have been identified in humans with myriad substrates including monosaccharides, short peptides, and ionic compounds.^15,16^ Since the abundance and localization of these transporters impacts the absorption of xenobiotic substrates in the small intestine, it is important to characterize SLC and ABC transporter expression in each region of the small intestine. For instance, previous mRNA studies in the adult small intestine observed regional differences in transporter levels as demonstrated by elevated mRNA transcript levels for ABCB1 (MDR1), SLC10A2 (ASBT), ABCG2 (BCRP), and SLC15A1 (PEPT1) in ileal compared to duodenal mucosa.^17,18^ Information on SLC and ABC transporter expression along the length of the neonatal and pediatric small intestine is also valuable as the data can inform decisions on dosing recommendations in pediatrics and will improve our understanding of how critical nutrients are absorbed throughout childhood development. Other groups have previously reported no change or a slight decrease in mRNA levels for ABCB1, ABCC2 (MRP2), and SLCO2B1 (OATP2B1) in infants and children compared to adults.^19^ The clinical relevance of determining mRNA transcript abundance is limited, however, as there may be poor correlation between mRNA levels and protein abundance for transporters in the small intestine.^20,21,22^

While there are reported to be more than 100 SLC and ABC transporter proteins expressed in the small intestine,^18,21,22^ fewer than 30 of these transporters have been quantified by targeted proteomics.^23^ The majority of previous studies quantify drug transporters despite the critical role of SLC and ABC transporters in nutrient absorption. Previous studies suggest drug transporters increase in abundance from proximal to distal small intestine mucosa in adults, though trends have not always been statistically significant.^18,21,22^ Transporter abundance data normalized to the enterocyte marker villin (VIL1) suggest no region-associated trend in neonates and pediatrics,^24^ while non-normalized results in neonates suggest that a few drug transporters including ABCG2 and SLC15A1 decrease from jejunum to ileum. By age, transporter (e.g., ABCB1 and ABCG2) abundance has been shown to increase significantly in jejunum and ileum from infancy to adulthood^24,26^ and in duodenum from young adults to geriatrics.^27^ To more thoroughly characterize nutrient and drug absorption in the small intestine, comprehensive quantification of SLC and ABC transporters by intestinal region in neonates, pediatrics, and adults is warranted.

Global proteomics provides an efficient approach to identify and quantify thousands of proteins in small intestine mucosa samples.^28^ Data-dependent acquisition (DDA) mass spectrometry (MS) is an untargeted proteomics approach that has been used to study drug-metabolizing enzymes and transporters in the adult small intestine.^23^ However, in complex samples such as intestine mucosa, DDA-MS is biased towards the identification of high-abundance proteins, necessitating extensive, resource-intensive fractionation of a representative sample to ensure deep proteome coverage.^29^ By comparison, data-independent acquisition (DIA) mass spectrometry is an efficient alternative global proteomics approach that overcomes many of the limitations of DDA-MS and can be used as a stand-alone technique to identify and quantify thousands of proteins in complex samples.^30^ We present here the DIA MS-based identification and relative quantification of proteins along the length of the small intestine in neonates, pediatrics, and adults, with emphasis on SLC and ABC transporters. To our knowledge, this study is the first to apply DIA-MS to elucidate the expression of thousands of small intestine proteins across intestinal region and age, and we highlight abundance trends for several understudied xenobiotic transporters in this critical organ.

## Materials and Methods

### Intestine sample collection

Small intestines from 18 donors (5 days – 34 years old, mixed sex, race: White or Black/African American) were procured from Novabiosis, Inc. (Morrisville, NC) or International Institute for the Advancement of Medicine (IIAM, Edison, NJ) (**Table 1**). Exclusion criteria are described in the Supplementary Material. Intestinal region (duodenum, jejunum, or ileum) was identified by gross anatomy during dissection as described in the Supplementary Methods. Mucosa was collected from each region of the small intestine by gently scraping with a razor blade and collected mucosa was snap frozen with liquid nitrogen, stored on dry ice for the duration of each intestine dissection, and then frozen at -80°C until preparation and trypsin digestion. All samples were de-identified prior to sample preparation and analysis.

**Table 1.** Number of mucosa samples by small intestine region for each age group.

| **Sample group** | **Duodenum** | **Jejunum** | **Ileum** | **Total** | **F/M** | **W/B** |
| --- | --- | --- | --- | --- | --- | --- |
| Neonate<br>(0-1 month) | 2 | 3 | 1 | 6 | 1/2 | 2/1 |
| Pediatric<br>(> 1 month - 8 years) | 5 | 6 | 7 | 18 | 2/5 | 4/2* |
| Adult<br>(18 - 34 years) | 3 | 7 | 7 | 17 | 4/4 | 7/1 |
| Total | 10 | 16 | 15 | 41 | 7/11 | 13/4* |

F = female, M = male, W = White, B = Black/African American, \*race not available for 1
pediatric donor

### Sample preparation

Mucosa was thawed and homogenized (5 m/s for 20 seconds) with an Omni Bead Ruptor 24 with Omni Cryo 24 in 2 mL tubes with 2.8 mm ceramic beads (Omni International, Revvity, Waltham, MA). The homogenization buffer was 500 µL 20 mM Tris (Fisher Scientific, Waltham, MA), 50 mM mannitol (Sigma Aldrich, St. Louis, MO), 1 mM EDTA (Sigma Aldrich, 99.999% purity, St. Louis, MO) at pH 7.4 (adjusted with HCl (Fisher Chemical, Waltham, MA)) with protease inhibitor (Roche cOmplete Tablets, EDTA-free, EASYpack, Basel, Switzerland). Protein concentration in mucosa homogenate was determined by the bicinchonic acid assay (Thermo Scientific Pierce, Waltham, MA) and samples were diluted to 0.5 mg/mL in 50 mM ammonium bicarbonate (Thermo Scientific, 99%, Waltham, MA) at pH 7.4 (pH adjusted using HCl). Sodium dodecyl sulfate (Fisher Scientific, Waltham, MA) was added to a final concentration of 1% and 1 µg of yeast enolase (Sigma Aldrich, St. Louis, MO) was prepared separately as a system suitability control. Samples were heated at 95°C for 5 mins with shaking (600 rpm) and allowed to cool to room temperature. Dithiothreitol (Thermo Scientific Pierce, Waltham, MA) was added to a final concentration of 10 mM and samples were incubated for 20 minutes at room temperature. Iodoacetamide (Thermo Scientific Pierce, Waltham, MA) was freshly prepared and added to a final concentration of 15 mM and samples were incubated for 20 minutes at room temperature in the dark.

MagReSyn beads (ReSyn Biosciences, Pretoria, South Africa) and samples were added to Kingfisher 96-well deep-well plates (Thermo Scientific, Waltham, MA) at a ratio of 250 µg beads:50 µg protein. Acetonitrile (Sigma Aldrich, St. Louis, MO) was added to bead-sample mixtures to a final concentration of 70% and incubated for 10 minutes at room temperature. The Kingfisher (Thermo Scientific Pierce, Waltham, MA) single-pot, solid-phase enhanced sample preparation method^31^ included ∼25 minutes of protein capture onto beads and 3 washes with 95% acetonitrile in water followed by 2 washes with 70% ethanol (Sigma Aldrich, St. Louis, MO) in water (∼2.5 minutes per wash) (Supplementary Material).

### Protein digestion

MS-grade trypsin (Thermo Scientific Pierce, Waltham, MA) was reconstituted in 50 mM glacial acetic acid (Fisher Chemical, Waltham, MA) in water at 0.5 mg/mL and diluted to 0.05 mg/mL with 50 mM ammonium bicarbonate in water, pH 7.4 immediately prior to digestion (20:1 protein:trypsin). Samples on beads were digested at 37°C for 16 hours with shaking at 1000 rpm on the Thermomixer. Once trypsin digestion was complete, samples were separated from beads on the Kingfisher and ice-cold MS-grade formic acid (Fisher Chemical, Bothell, WA) was added to samples to a final concentration of 5%. Samples were centrifuged at 14,000 *g* for 15 mins at 4°C to pellet remaining magnetic beads, and supernatant was dried by Speed Vac (Savant SPD1030 Integrated Vacuum Concentrator (Thermo Scientific, Waltham, MA)) at 45°C and a target pressure of 14 mTorr with dried samples stored at -80°C until liquid chromatography-tandem mass spectrometry (LC-MS/MS) analysis.

### Peptide sample analysis by DIA-MS

Peptide samples were reconstituted (0.5 mg/mL, 1 µg injected on-column) in the following solution: 2% acetonitrile (Optima LC/MS grade, Fisher Chemical, Bothell, WA), 0.1% formic acid, 25 fmol/µL Pierce peptide retention time calibration mixture (Thermo Scientific Pierce, Waltham, MA) in water. A Thermo Scientific Vanquish Neo UHPLC was used with a Thermo Scientific PepMap Neo C18 trap column (5 µm, 300 µm x 5 mm, Waltham, MA), a Bruker PepSep Max C18 column (10 cm x 150 µm, 1.5 µm, Billerica, MA), and a FossilionTech nanoESI emitter (Sharp Singularity Classic Ref 20-05, 20 µm, 5 cm, Madrid, Spain). Mobile phase A was 0.1% formic acid in water (Optima, Fisher Chemical, Bothell, WA) and mobile phase B was 0.1% formic acid (Optima, Fisher Chemical, Bothell, WA) in 80% acetonitrile (Optima, Fisher Chemical, Bothell, WA) in water. The column temperature was 35°C and the injection volume was 2 µL. The autosampler was set at 4°C. The liquid chromatography gradient was as follows: 4-6% B at 1.3 µL/min (0-0.7 min), 6-10% B at 1.3 µL/min (0.7-1 min), 10%- 50% B at 0.8 µL/min (1-26 mins), 50-99% B at 0.8-1.3 µL/min (26-26.5 mins), 99% B at 1.3 µL/min (26.5-30 mins). A Thermo Scientific Orbitrap Astral was used for DIA-MS. Individual peptide samples were analyzed with wide-window DIA runs (expected precursor peak width of 25 seconds and charge of 3, positive polarity, normalized collision energy (NCE) of 27, ion injection time of 3.5 ms, automatic gain control target of 1,000,000) of 8 m/z windows (with overlapping by 4 m/z) spanning 400-1004 m/z, with window placement optimized with Skyline.^32^ A MS scan from 390-1010 m/z for precursor spectra (expected precursor peak width of 25 seconds and charge of 3, positive polarity, 60,000 resolution, ion injection time of 3.5 ms, automatic gain control target of 1,000,000) was interspersed every 75 MS/MS spectra.

### DIA data analysis

DIA spectra were demultiplexed with ProteoWizard (peak picking, 10 ppm accuracy) (version 3.0.24075)^33^ and mzML files were searched against a Prosit-^34^ or Koina-predicted^35^ spectral library with EncyclopeDIA (version 2.12.30).^30^ The spectral library was based on a combined human proteome FASTA (UP000005640, downloaded on December 08, 2023) and yeast ENO1 FASTA (downloaded on March 04, 2024) from Uniprot. Prosit or Koina were run with tryptic digestion with 1 missed cleavage, a peptide precursor range from 390-1010 m/z and charge of +2 or +3, default NCE of 33 adjusted for DIA, default precursor charge of +3, the intensity model Prosit_2020_intensity_HCD, and the indexed retention time model Prosit_2019_irt at a NCE of 33. EncyclopeDIA was configured with default settings including normal target/decoy search; tryptic peptides; CID/HCD fragmentation with b/y ions; 10 ppm precursor, fragment, and library tolerance; and ≥ 3 quantitative ions. Percolator (version 3-01)^36^ was used to control the false discovery rate at 1% at the peptide and protein levels.

Peptide results were visualized with Skyline and a background proteome was generated based on the combined human proteome and yeast ENO1 FASTA. Results were filtered in Skyline to include only peptides with ≥ 3 b/y fragment ions and only proteins with ≥ 1 unique peptide based on the background proteome. Peptides were manually curated in Skyline to remove unlikely peptide spectrum matches. Results were exported from Skyline for additional analysis with MSstats.^37^ Fragment peak areas were log_2_ transformed, median-normalized, and summarized with Tukey’s median polish using MSstats to obtain protein peak areas. Fragment peak areas < 1 were censored. Protein peak areas were averaged across the two experimental replicates per sample to obtain one protein peak area for each sample.

The average number of proteins and SLC/ABC transporters identified across the two experimental replicates per sample were plotted with GraphPad Prism 11 (version 11.0.1). Venn diagrams comparing the average number of proteins identified in ≥ 1 sample (in both experimental replicates) per condition after the filtering criteria above were created with the ggVennDiagram package in R. The log_2_-transformed average protein peak area was used for dynamic range plots of proteins identified in every experimental replicate for every sample by type (e.g., every sample per age group or small intestinal region). Select nutrient and drug transporters were highlighted in the dynamic range plots. Principal component analysis (PCA) of proteins identified in every experimental replicate of all samples was conducted. Biplots of principal components 1 and 2 were color-coded by sample metadata (e.g., age group).

Differential protein expression was determined by Student’s t-tests of average protein peak area with multiple testing corrected by the Benjamini-Hochberg method (performed with the limma package in R). Statistically significant differential protein expression was based on an adjusted p-value < 0.05 and a log_2_-transformed fold change ≥ ± 2. Volcano plots display proteins compared across pairs of age groups controlling for small intestinal region (e.g., pediatric jejunum compared to adult jejunum) or small intestinal regions controlling for age (e.g., pediatric duodenum compared to pediatric jejunum). Small intestinal regions were not compared in neonates due to the small number of samples available.

### Data records

The complete datasets for the DIA-MS studies are available via Panorama Public^38^ (https://panoramaweb.org/uwtec_gi_mucosa_transporters.url) and were assigned the ProteomeXchange ^39^ ID: PXD079663 (https://doi.org/10.6069/jb88-rg90). Experimental data are provided at multiple levels of processing, from raw DIA-MS data to normalized peptide- and protein-level quantities suitable for downstream analysis.

## Results

### Donor demographics and number of proteins

The DIA-MS workflow used in this study allowed for the identification and relative quantification of thousands of proteins in small intestine mucosa samples (**Table 1**; donor demographics in **Table S1**). On average 5,229 proteins (standard deviation (sd): 88 proteins) were identified in all samples, including 87 SLC transporters (sd: 2) and 18 ABC transporters (sd: 0.3). The number of proteins and SLC or ABC transporters did not change notably with small intestinal region or age group (**Table 2**).

**Table 2.** Number of protein and SLC/ABC transporters by small intestinal region and age group.

|  | Small intestine region |  |  | Age group |  |  |
| --- | --- | --- | --- | --- | --- | --- |
|  | Duodenum | Jejunum | Ileum | Neonate<br>(0-1 month) | Pediatric<br>(1 month-8 years) | Adult<br>(18 years-34 years) |
| n | 10 | 16 | 15 | 6 | 18 | 17 |
| Total proteins | 5,278 (161) | 5,198 (350) | 5,194 (325) | 5,284 (244) | 5,276 (267) | 5,127 (338) |
| SLC<br>transporters | 86 (6) | 85 (8) | 88 (5) | 88 (5) | 88 (7) | 84 (7) |
| ABC<br>transporters | 19 (0.8) | 18 (1) | 19 (1) | 19 (1) | 19 (1) | 18 (1) |

Mean (standard deviation)

### Sample clustering and unique proteins

Principal component analysis (PCA) indicated mucosa samples clustered by small intestinal region more strongly than by age group (**Figure S1**). The majority of duodenum and jejunum samples clustered apart from ileum samples (**Figure S1A**). Many neonate samples clustered apart from adult samples, but pediatric samples overlapped with neonate and adult samples (**Figure S1B**). Samples did not cluster strongly by donor sex (**Figure S1C**).

The majority of identified proteins were observed in all small intestinal regions and age groups (**Figure 1A** and **1B**). Approximately 5,500 proteins were shared among small intestinal regions or age groups, and fewer than 1% of proteins were unique to an intestinal region or age group. Two SLC transporters appeared to be specific to the ileum, SLC10A2 and SLC6A20, and SLC7A5 appeared to be unique to pediatric samples in all small intestinal regions. When comparing protein expression by intestinal region within neonate, pediatric, or adult sample sets, the results were similar to those observed when all ages were combined together (**Figure 1C-1E**). Approximately 5,420 proteins were expressed in all intestinal regions within each age group, and the same two transporters (SLC10A2 and SLC6A20) were unique to ileum in all age groups.

**Figure 1.**
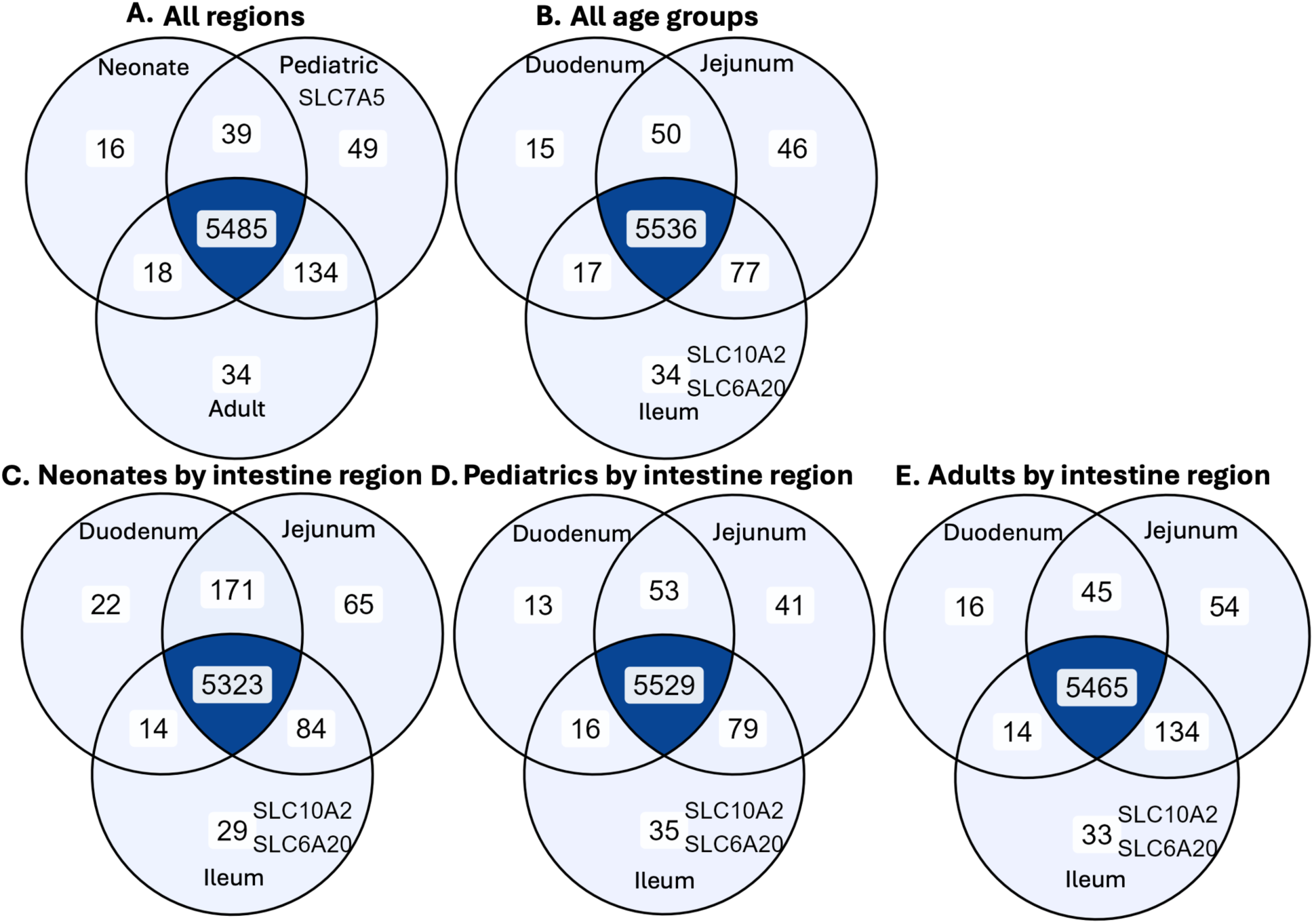
Total number of shared and unique proteins by small intestinal region and/or age group. Unique SLC/ABC transporters are listed. All proteins were present in both experimental replicates of ≥ 1 sample per condition.

### Small intestine markers and dynamic range plots

The expression of the small intestine marker proteins VIL1 and MUC2 increased from proximal to distal small intestine (**Figure 2**). When comparing ileum to duodenum in pediatrics and adults, VIL1 had a log_2_ fold change of 0.77 (p = 0.02) and MUC2 had a log_2_ fold change of 1.85 (p = 0.0002). In contrast, other small intestine marker proteins (ATP1A1, ALPI, and SI) had no significant abundance trends with region or age group (**Figures S2 and S3**). The relative abundance of approximately half of a selection of SLC or ABC transporters was similar across small intestinal region in neonates, pediatrics, and adults (**Figure 3**). Among a group of select SLC and ABC transporters, SLC5A1 (SGLT1) and SLC6A19 had relatively high abundance while SLCO2B1 had relatively low abundance. ABCB1 and SLC15A1 generally increased in abundance from duodenum to ileum in samples from all age groups. While the relative abundance of SLC2A5 (GLUT5) was higher in pediatric and adult duodenum compared to neonate duodenum, most of the selected SLC or ABC transporters did not display strong age-associated trends in relative abundance in jejunum or ileum (**Figure 3**).

**Figure 2.**
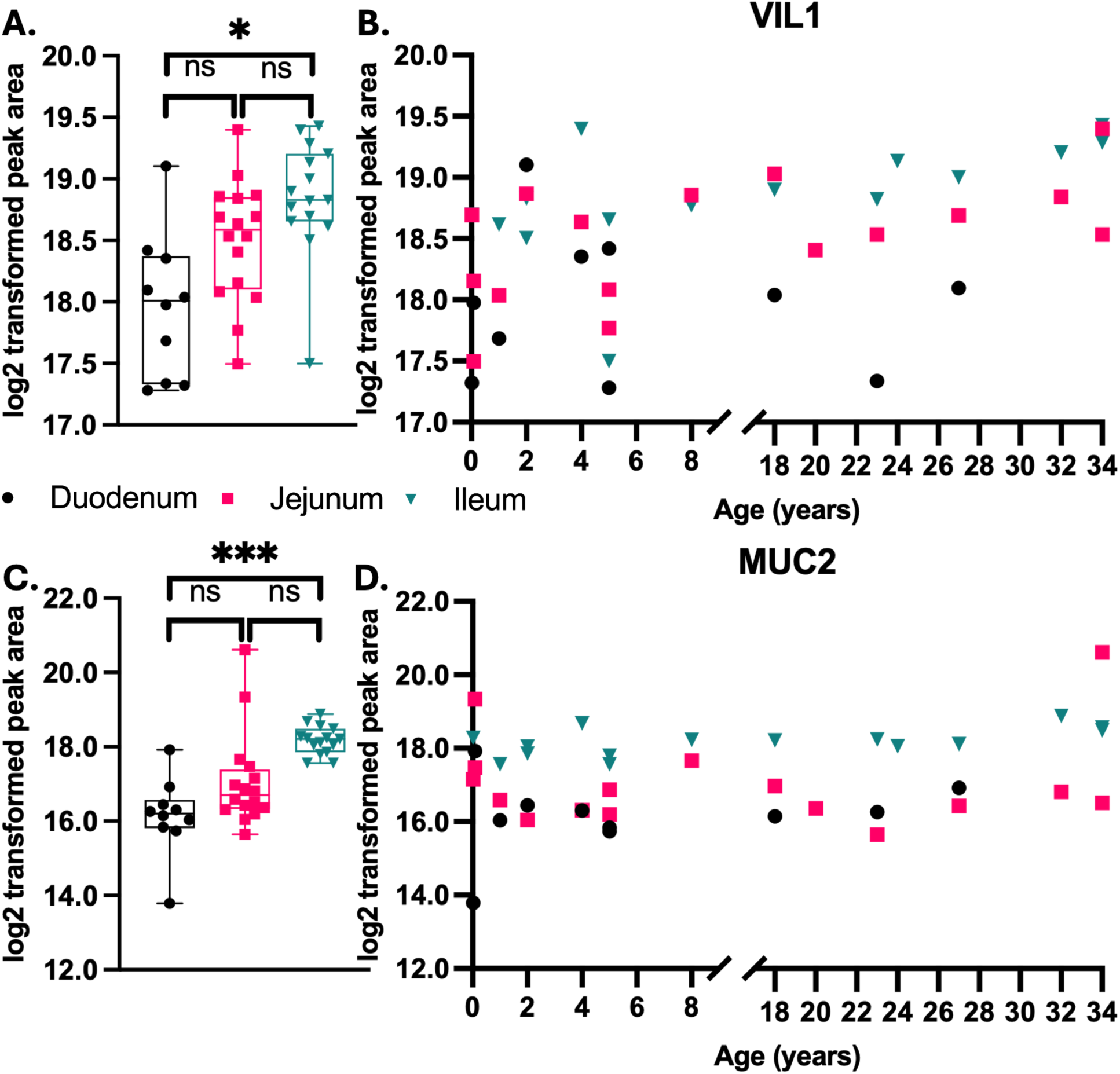
VIL1 and MUC2 abundance by intestinal region and age. VIL1 peak area increases by small intestinal region (A) but not with age (B), which was also observed for MUC2 by region (C) and with age (D). n = 10 duodenum, n = 16 jejunum, n = 15 ileum; n = 1 for all ages other than n = 2 for 1 month old, 2 years old, 5 years old; t-tests for jejunum/duodenum: n = 5,831 proteins, for ileum/jejunum: n = 5,846 proteins, for ileum/duodenum: n = 5,824 proteins, with multiple testing corrected by Benjamini-Hochberg method, ns = not significant, *adjusted p < 0.05, ***adjusted p < 0.001

**Figure 3.**
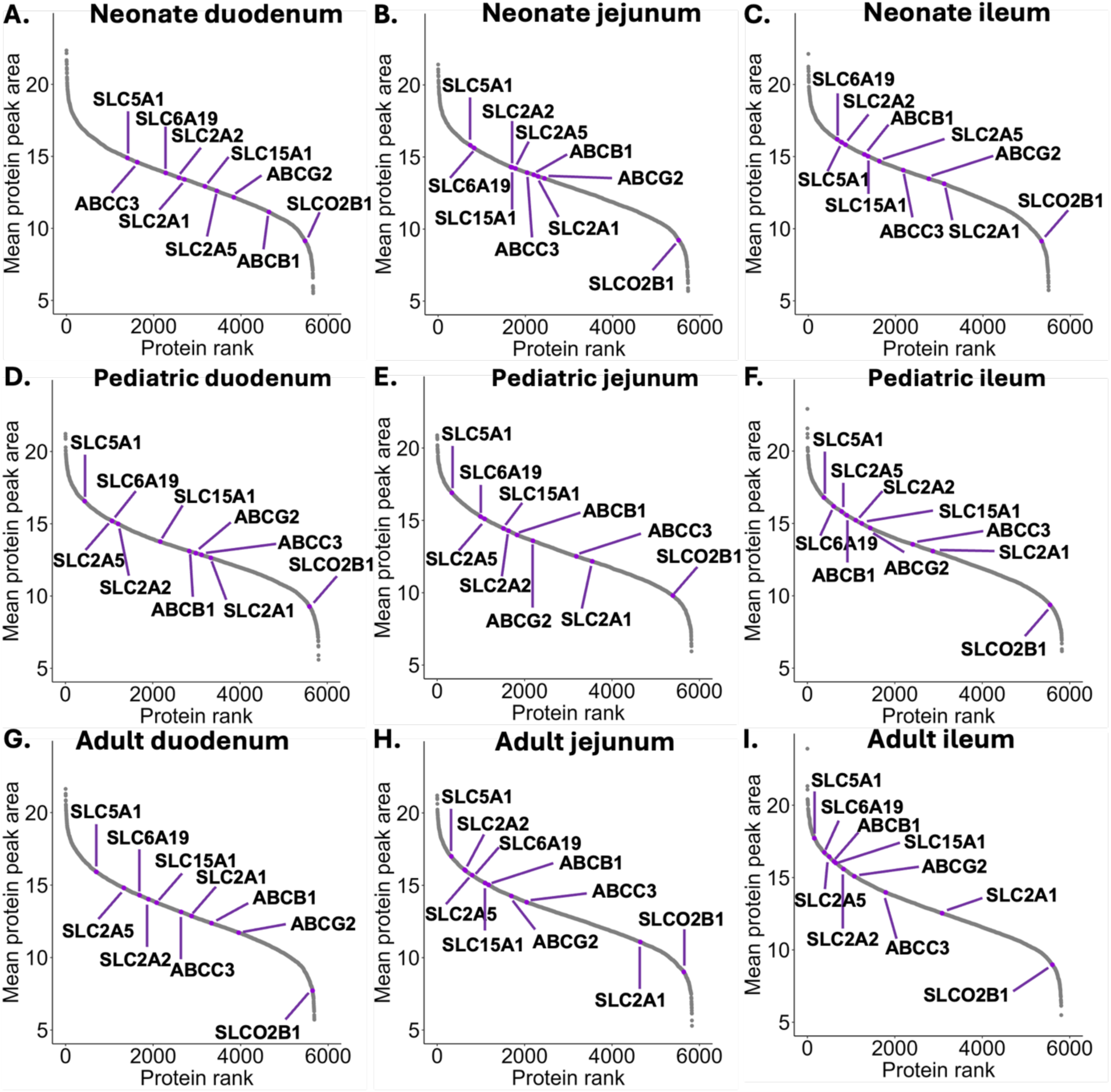
Relative abundance of select SLC/ABC transporters by intestinal region for each age group. Relative abundances are shown for neonates (duodenum n = 5,654, jejunum n = 5,727, ileum n = 5,503) (A-C), pediatrics (duodenum n = 5,800, jejunum n = 5,818, ileum n = 5,821) (D-F), adults (duodenum n = 5,684, jejunum n = 5,829, ileum n = 5,805) (G-I).

### Region- and age-associated differences in transporter expression in the small intestine

Several SLC or ABC transporters were differentially expressed by small intestinal region. Although pediatric duodenum and jejunum had similar proteomes (**Figure 4A**), in adults 108 proteins in these two regions were differentially expressed, including 10 SLC or ABC transporters (**Figure 4D**, **Table 4**). When jejunum and ileum were compared, 90 proteins were differentially expressed in pediatrics including 4 SLC or ABC transporters (**Figure 4B and Table 4**), while 103 proteins were differentially expressed in adults including 8 SLC or ABC transporters (**Figure 4E and Table 4**). When duodenum and ileum were compared, 46 proteins were differentially expressed in pediatrics including 4 SLC or ABC transporters (**Figure 4C and Table 3**), and 271 proteins were differentially expressed in adults including 20 SLC or ABC transporters (**Figure 4F and Table 4**). A few SLC/ABC transporters were differentially expressed by intestinal region only in pediatrics or in adults. SLC26A3 and SLC51A (OSTα) were differentially expressed by intestinal region in pediatrics only (**Table 3**), while 21 SLC or ABC transporters were differentially expressed by intestinal region in adults only (**Table 4**). For example, SLC6A19 increased in abundance from duodenum to ileum in adults (log_2_ fold change of 2.5, p = 0.01) but not significantly in pediatrics (log_2_ fold change of 0.99, p = 0.61) (**Figure 5A**). Similarly, ABCG2 had higher expression in adult ileum than in duodenum (log_2_ fold change of 3.1, p = 0.0004) and in adult jejunum compared to duodenum (log_2_ fold change of 2.3, p = 0.02), but showed no significant change in pediatric ileum compared to duodenum (log_2_ fold change of 1.7, p = 0.18) or pediatric jejunum compared to duodenum (log_2_ fold change of 0.6, p = 0.72) (**Figure 5D**). ABCC2 also had increased abundance in adult ileum compared to duodenum (log_2_ fold change of 2.0, p = 0.03) but was not significantly different in pediatrics by intestinal region (ileum/duodenum log_2_ fold change of 1.2, p = 0.75) (**Figure 5E**).

**Figure 4.**
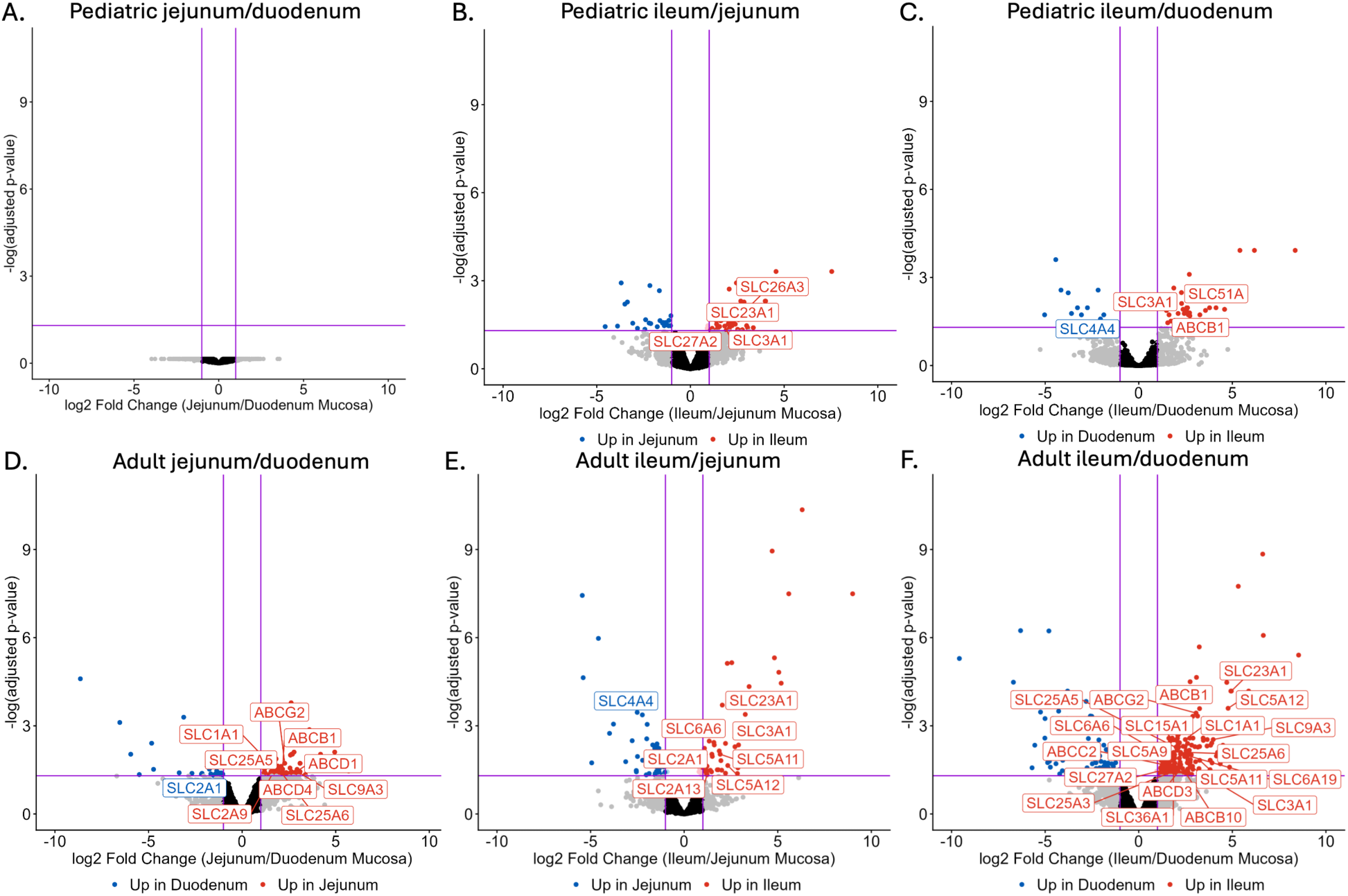
Differentially expressed SLC/ABC transporters by intestinal region. Volcano plots are shown for all proteins identified in pediatrics (A-C) and adults (D-F) with a subset of significantly differentially expressed SLC and ABC transporters labeled. n = 5 pediatric duodenum, n = 6 pediatric jejunum, n = 7 pediatric ileum, n = 3 adult duodenum, n = 7 adult jejunum, n = 7 adult ileum; t-tests with multiple testing corrected by Benjamini-Hochberg method; pediatric duodenum versus jejunum n = 5,762 proteins, pediatric jejunum versus ileum n = 5,784 proteins, pediatric duodenum versus ileum n = 5,759 proteins, adult duodenum versus jejunum n = 5,657 proteins, pediatric jejunum versus ileum n = 5,771 proteins, pediatric duodenum versus ileum n = 5,648 proteins

**Figure 5.**
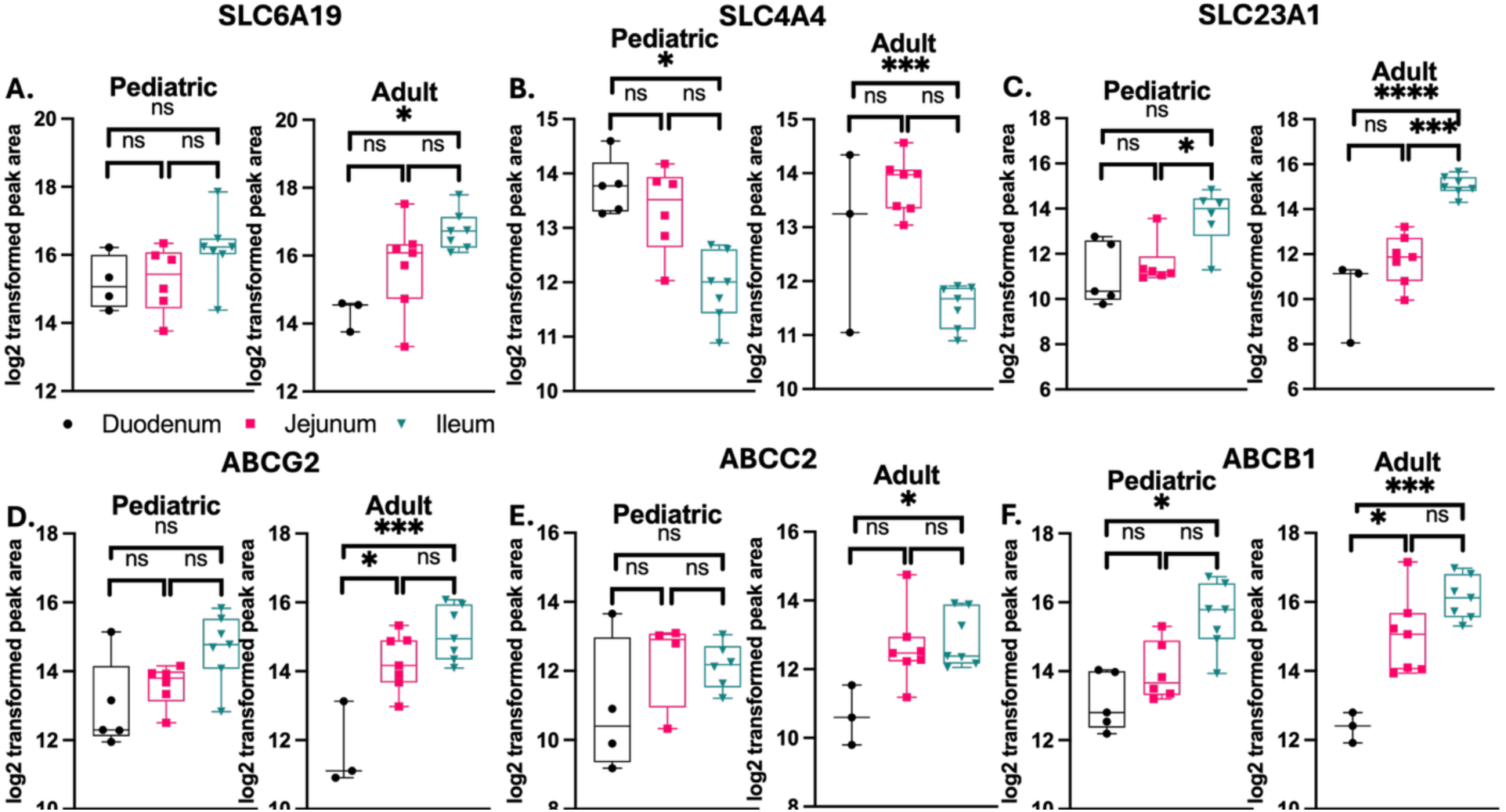
Select SLC/ABC transporter abundance by intestinal region and age. n = 10 duodenum, n = 16 jejunum, n = 15 ileum; n = 1 for all ages other than n = 2 for 1 month old, 2 years old, 5 years old; t-tests for jejunum/duodenum: n = 5,831 proteins, for ileum/jejunum: n = 5,846 proteins, for ileum/duodenum: n = 5,824 proteins, with multiple testing corrected by Benjamini-Hochberg method, ns = not significant, *adjusted p < 0.05, ***adjusted p < 0.001, ****adjusted p < 0.0001

**Table 3.** Differentially expressed SLC/ABC transporters by intestinal region in pediatrics.

| Comparison | Gene | Log<sub>2</sub> fold change | Adjusted p-value | Putative function |
| --- | --- | --- | --- | --- |
| i-j | SLC26A3 | 3.14 | 0.008 | Cl<sup>-</sup> and bicarbonate exchange (2:1) |
| i-j | SLC23A1 | 2.59 | 0.02 | Na<sup>+</sup> dependent uptake of vitamin C |
| i-j | SLC27A2 | 1.49 | 0.04 | long-chain fatty acid uptake |
| i-j, i-d | SLC3A1 | 1.83, 2.37 | 0.04, 0.01 | Cationic amino acid uptake and neutral amino acid efflux |
| i-d | SLC51A* | 2.61 | 0.01 | basolateral bile acid and steroid efflux |
| i-d | SLC4A4 | -1.87 | 0.02 | basolateral Na<sup>+</sup> and bicarbonate transport |
| i-d | ABCB1* | 2.43 | 0.02 | xenobiotic efflux |

j-d: jejunum/duodenum, i-j: ileum/jejunum, i-d: ileum/duodenum, \*has quantitative peptides

**Table 4.** Differentially expressed SLC/ABC transporters by intestinal region in adults.

| Comparison | Gene | Log<sub>2</sub> fold change | Adjusted p-value | Putative function |
| --- | --- | --- | --- | --- |
| j-d | SLC2A9 | 1.57 | 0.049 | urate uptake |
| j-d | ABCD1 | 2.83 | 0.04 | peroxisomal very long chain fatty acid uptake |
| j-d | ABCD4 | 1.93 | 0.048 | lysosomal vitamin B12 efflux |
| i-j | SLC4A4 | -2.23 | 4.2e-04 | basolateral Na<sup>+</sup> and bicarbonate transport |
| i-j | SLC2A13 | 1.22 | 0.046 | H<sup>+</sup> dependent myo-inositol transport |
| j-d, i-j | SLC2A1 | -1.67, 1.50 | 0.04, 0.01 | glucose uptake |
| j-d, i-d | SLC25A5* | 1.84, 2.08 | 0.04, 5.9e-03 | mitochondrial ATP efflux and ADP uptake |
| j-d, i-d | SLC1A1 | 1.96, 2.49 | 0.04, 2.9e-03 | Na<sup>+</sup> dependent glutamate or aspartate uptake with 1 H<sup>+</sup> and efflux of 1 K<sup>+</sup> |
| j-d, i-d | SLC9A3 | 3.07, 4.00 | 0.04, 2.9e-03 | H<sup>+</sup> efflux, Na<sup>+</sup> uptake |
| j-d, i-d | SLC25A6* | 2.17, 2.58 | 0.045, 8.0e-03 | mitochondrial ATP efflux and ADP uptake |
| j-d, i-d | ABCB1* | 2.16, 3.25 | 0.03, 2.6e-04 | xenobiotic efflux |
| j-d, i-d | ABCG2* | 2.28, 3.06 | 0.02, 3.7e-04 | xenobiotic, porphyrin, heme, steroid efflux |
| i-j, i-d | SLC23A1 | 3.26, 4.95 | 4.1e-04, 6.6e-05 | Na<sup>+</sup> dependent uptake of vitamin C |
| i-j, i-d | SLC5A12* | 2.33, 4.77 | 0.02, 2.5e-04 | low-affinity Na<sup>+</sup> dependent monocarboxylate uptake |
| i-j, i-d | SLC5A11 | 1.92, 3.26 | 9.3e-03, 0.01 | Na<sup>+</sup> dependent myo-inositol uptake |
| i-j, i-d | SLC6A6 | 1.59, 1.86 | 3.7e-03, 0.02 | Na<sup>+</sup> and Cl<sup>-</sup> dependent transport of taurine |
| i-j, i-d | SLC3A1* | 2.91, 2.82 | 4.5e-03, 0.03 | Cationic amino acid uptake and neutral amino acid efflux |
| i-d | SLC5A9 | 1.86 | 0.02 | Na<sup>+</sup> dependent mannose uptake |
| i-d | SLC15A1* | 2.10 | 4.3e-03 | H<sup>+</sup> dependent di- or tri-peptide uptake |
| i-d | SLC6A19* | 2.48 | 9.8e-03 | Na<sup>+</sup> dependent neutral amino acid uptake |
| i-d | SLC27A2 | 1.46 | 0.03 | long-chain fatty acid uptake |
| i-d | SLC25A3* | 1.55 | 0.04 | mitochondrial phosphate (H<sup>+</sup> dependent) and copper uptake |
| i-d | SLC36A1* | 1.86 | 0.04 | H<sup>+</sup> dependent small amino acid transport |
| i-d | ABCC2 | 1.95 | 0.03 | xenobiotic, glucuronide/sulfate/glutathione conjugate efflux |
| i-d | ABCB10 | 2.47 | 0.04 | mitochondrial biliverdin efflux |
| i-d | ABCD3* | 2.05 | 0.04 | peroxisomal bile acid and long chain fatty acid uptake |

j-d: jejunum/duodenum, i-j: ileum/jejunum, i-d: ileum/duodenum, \*has quantitative peptides

Nevertheless, several transporters were differentially expressed by intestinal region in both pediatrics and adults (**Table 3 and 4**). For instance, SLC4A4, a bicarbonate transporter that maintains small intestine lumen pH,^40^ displayed decreasing abundance from duodenum to ileum in pediatrics (log_2_ fold change of 1.9, p = 0.02) and from jejunum to ileum in adults (log_2_ fold change of 2.2, p = 0.0004) (**Figure 5B**). The ascorbic acid uptake transporter SLC23A1 showed the opposite trend, increasing from jejunum to ileum in pediatrics (log_2_ fold change of 2.6, p = 0.02) and adults (log_2_ fold change of 3.3, p = 0.0004) and from duodenum to ileum in adults only (log_2_ fold change of 5.0, p = 0.00007) (**Figure 5C**). Similarly, ABCB1 increased from duodenum to ileum in pediatrics (log_2_ fold change of 2.4, p = 0.02) and adults (log_2_ fold change of 3.3, p = 0.0003) and from duodenum to jejunum in adults only (log_2_ fold change of 2.2, p = 0.03) (**Figure 5F**). Of the 29 SLC or ABC transporters that were differentially expressed by either intestinal region or age group, 12 (41%) had at least one quantitative peptide based on the matched-matrix calibration curve, which increases confidence that the observed regional differences in protein peak area for these transporters corresponded to differences in protein abundance.

In contrast, few SLC or ABC transporters were differentially expressed when age groups were compared. When neonatal and pediatric mucosa were compared, only 12 proteins were significantly differentially expressed, and none of the proteins were transporters (**Figure S4G**). Similarly, for pediatric compared to adult mucosa 36 proteins were differentially expressed, including SLC13A2, which was higher in adults than in pediatrics (log2 fold change of 1.6, p = 0.03) (**Figure S4H**). While 17 proteins were differentially expressed between neonatal and adult mucosa, SLC13A2 was the only transporter that was higher in adults than in neonates (log2 fold change of 3.7, p = 0.006) (**Figure S4I**).

## Discussion

In this study, DIA-MS was used to characterize the abundance of nutrient and drug transporters along the length of the small intestine in neonates, pediatrics, and adults. The majority of previous studies of xenobiotic transporter abundance in the small intestine focused on ∼30 SLC and ABC transporters,^18,24,25^ overlooking ∼80 additional transporters believed to have important roles in nutrient and drug uptake and efflux in the intestine^23^. To address this gap in knowledge, global proteomics is increasingly being used to study transporter expression in the small intestine.^23^ However, we are aware of only one previous DIA MS-based analysis of adult small intestine, which focused primarily on intestinal tissue biopsies (i.e., samples were not limited to mucosa) from patients with inflammatory bowel disease, which may not reflect healthy intestinal mucosa.^41^ In the present study, mucosa from small intestine organ donors with no history of gastrointestinal trauma or disease was collected, enabling the investigation of SLC and ABC transporter expression along the small intestine in neonates, pediatrics, and adults. To our knowledge, this study is the first to use DIA-MS to elucidate transporter abundance trends with intestinal region in neonates and pediatrics. Consistent with the deep proteome coverage associated with DIA-MS, an average of 5,229 proteins including ∼90 SLC and ∼20 ABC transporters were identified across all mucosa samples. The total number of proteins as well as SLC and ABC transporters identified in this study was similar to previously published results using DDA-MS with adult jejunum and ileum mucosa.^23^

Small intestine marker proteins were analyzed to confirm the presence of epithelial cells in all samples and identify any trends in marker protein abundance by intestinal region. There was an increase in VIL1 abundance from duodenum to ileum (log_2_ fold change of 0.77, p = 0.02) and a non-significant upward trend with age (in adults versus neonates, log_2_ fold change of 0.62, p = 0.77), matching results previously observed in cryopreserved adult duodenum and ileum mucosa^42^ and in neonate and adult jejunum and ileum.^24^ Several previous studies have normalized transporter abundance to VIL1 to account for differences in enterocyte count among samples,^23,24,42^ with the assumption that VIL1 abundance does not change with small intestinal region or age. However, our results highlight the potential confounding effect of VIL1 normalization of protein abundance. MUC2 would also be a poor candidate for normalization, as its abundance increased significantly along the small intestine in this study (duodenum to ileum log_2_ fold change of 1.85, p = 0.0002), consistent with previous reports in animal models^43^ and in human jejunum and ileum.^23^ Other marker proteins such as ATP1A1, ALPI, and SI may be appropriate for normalization since they displayed no trends with intestinal region or age in this study, though others have reported a significant increase in SI from duodenum to ileum^42^ or decreasing expression of ATP1A1, ALPI, and SI from jejunum to ileum.^23^ These contradictory results reflect the need for caution when normalizing to small intestine marker proteins.

The DIA-MS workflow employed in this study enabled the identification of significant abundance differences for both previously studied and less well-characterized transporters along the length of the small intestine in different age groups. This study identified 105 SLC and ABC transporters across all samples in experimental triplicate, whereas previous targeted proteomics studies quantified only up to 11 of these transporters.^18,23,24,26^ In agreement with previous reports,^18,44^ this study observed that ABCB1 had increasing peak area from proximal to distal small intestine (in pediatric duodenum to ileum log_2_ fold change of 2.4, p = 0.02; in adult duodenum to ileum log_2_ fold change of 3.3, p = 0.0003), and both SLC15A1 (log_2_ fold change of 2.1, p = 0.004) and ABCG2 (log_2_ fold change of 3.1, p = 0.0004) had significantly higher peak area in ileum than in duodenum in adults but not in pediatrics (SLC15A1 log_2_ fold change of 1.2, p = 0.44; ABCG2 log_2_ fold change of 1.7, p = 0.18). Many of the additional transporters that could be quantified in our data are infrequently studied nutrient transporters with poorly characterized abundance along the small intestine, highlighting the value of the global proteomics approach used here. For instance, SLC4A4 had higher peak area in duodenum and jejunum than in ileum (pediatric duodenum to ileum log_2_ fold change of -1.9, p = 0.02; adult jejunum to ileum log_2_ fold change of -2.2, p = 0.0004), but to our knowledge its intestinal regional expression has not been reported previously. Although SLC7A5 appeared to be unique to pediatric small intestine in our results, this amino acid transporter heterodimerizes with SLC3A2 for function, and SLC3A2 was not unique to pediatrics. Thus, it is unclear whether SLC7A5 is truly unique to pediatrics and the observed result may be due to the low abundance of this protein in all age groups.^45^ While 31 of the 105 SLC and ABC transporters in this study were previously quantified in adult jejunum and ileum by others using DDA-MS, the expression of many understudied xenobiotic transporters is now available in all small intestine regions for neonates, pediatrics, and adults.^23^

Interestingly, while SLC10A2 is known to be localized predominantly in the ileum,^18,24^ our data suggest that SLC6A20 may also be uniquely expressed in ileum mucosa. Our findings are consistent with results from previous targeted proteomics studies, in which SLC10A2 was unique to adult ileum^24^ or expressed at a higher level in pediatric and adult ileum compared to jejunum.^18^ The ileum-specific localization of SLC10A2 in this study may be attributable to its low abundance in duodenum and jejunum samples making this transporter undetectable by DIA-MS in proximal regions. Previous investigation of SLC6A20, which facilitates the absorption of proline and N-methylated amino acids, included immunolocalization studies in the adult ileum^46^ and transport kinetics studies in fetal jejunum and ileum.^47^ Notably, quantifiable SLC6A20 activity in fetal small intestine^47^ supports our finding that this transporter is expressed in neonates and pediatrics. Moving forward, the selective ileal expression of SLC6A20 should be confirmed with orthogonal approaches such as targeted proteomics and/or immunohistochemistry.

Several SLC and ABC transporters were differentially expressed by small intestinal region within each age group. We identified 7 SLC/ABC transporters that were significantly differentially expressed by small intestinal region in pediatrics and 26 in adults. Only 5 of the transporters (SLC4A4, SLC23A1, SLC3A1, SLC27A2, ABCB1) were differentially expressed by intestinal region in both age groups. Potential reasons for the discrepancy in differentially expressed transporters by intestinal region in pediatrics compared to adults may include the limited number of donors in each age group (7 pediatrics and 8 adults) and missing intestinal regions for several donors (e.g., duodenum was available for only 3 of the 8 adult donors) (**Table 1**). Confirmatory targeted proteomics studies are warranted, as 17 SLC and ABC transporters in this study were differentially expressed by intestinal region but lacked quantitative peptides based on the matched-matrix calibration curve.

We acknowledge that in addition to the new findings reported in this study, several SLC and ABC transporters previously reported to have differential expression by intestinal region and/or age displayed no significant trends in our results. Of the 23 SLC and ABC transporters that were differentially expressed by small intestinal region in this study, only 4 (SLC15A1, ABCB1, ABCC2, ABCG2) were previously reported to have significant differential expression based on targeted proteomics.^18^ Our study did not find intestinal region-associated abundance for SLC16A1, SLC22A1, SLCO2B1, or ABCC3, contrary to a previous report in adults.^18^ The one transporter that we identified as more abundant in pediatrics and adults than in neonates, SLC13A2, has not been previously reported as such. In comparison, others have identified 4 SLC or ABC transporters (SLC15A1, ABCB1, ABCC2, ABCG2) that differ significantly with age based on targeted proteomics,^24^ but we did not observe significant age-associated trends for those transporters. The discrepancy between our study and previous studies may be attributable to relatively low statistical power in this study due to the small number of donors in each age group. Furthermore, certain small intestinal regions were not available for a subset of donors (e.g., duodenum was missing from donors also donating the pancreas) in this study, whereas all regions were available for adult donors in other studies,^18^ leading to greater statistical power. Moreover, DIA-MS has lower sensitivity compared to targeted proteomics and protein abundance in this study was determined by summarizing peptide peak areas instead of using a specific, quantitative peptide for each protein. While 9 transporters studied previously by targeted proteomics in small intestine mucosa were not identified in our data, only SLC22A1 (OCT1) was detected and quantified in multiple previous studies.^18,23,24,26^ The absence of SLC22A1 in our data set may be due to limitations associated with DIA-MS. Since DIA-MS fragments all peptide precursors within a given m/z range for peptide identification, mass spectrometer parameters are not optimized for specific transporter peptides, and may provide lower sensitivity than optimized targeted proteomics methods.^24,28^ Protein identification in this study is based on unique peptides that generate detectable b and y fragment ions at the collision energy used for each m/z range, instead of based on a curated list of quantitative peptides. Follow-up targeted proteomics is necessary to confirm the absence of SLC22A1 in our mucosa samples.

In conclusion, this study used a DIA MS-only workflow to identify and quantify drug transporters and less well-characterized nutrient transporters along the length of the small intestine in neonates, pediatrics, and adults. We identified several differentially expressed SLC and ABC transporters by small intestinal region in pediatrics and adults, and the results reported here will serve as a valuable reference for future prediction of xenobiotic and nutrient absorption and efflux across age.

## Abbreviations

ABC: ATP-binding cassette
B: Black/African American
d: duodenum
DDA: data-dependent acquisition
DIA: data-independent acquisition
F: female
MS: mass spectrometry
i: ileum
IIAM: International Institute for the Advancement of Medicine
j: jejunum
LC-MS/MS: liquid chromatography tandem mass spectrometry
M: male
NCE: normalized collision energy
ns: not significant
PCA: principal component analysis
sd: standard deviation
SLC: solute carrier
W: White

## Declaration of generative AI use

The authors used Microsoft CoPilot to assist with generating R code with the sole purpose of formatting data prior to analysis with published R packages. After using this service, the authors reviewed the data and output as needed and take full responsibility for the content of the published article.

## Acknowledgments

We would like to thank Dr. Chris Arian, Eimear O’Mahony, Brooke Chalker, Dr. Tim Tsang, Katharine Staudinger, and Yiqing Wang for assistance processing intestine tissue. Additionally, we thank Dr. Alex Zelter for assistance with the Kingfisher sample preparation method and LC-MS/MS method.

## Data availability

All data supporting the findings of this study are available within this paper and its supplementary material. Proteomic data presented in the current work are available via Panorama Public^38^ (https://panoramaweb.org/uwtec_gi_mucosa_transporters.url) and were assigned the ProteomeXchange ^39^ ID: PXD079663 (https://doi.org/10.6069/jb88-rg90).

## Authorship contributions

**VS** contributed to investigation, data curation, formal analysis, visualization, writing – original draft; **KW** contributed to investigation; **EJK** contributed to funding acquisition, resources, writing – reviewing and editing ; **SLMA** contributed to conceptualization, funding acquisition, methodology, project administration, supervision, writing – reviewing and editing.

## References

(1) Goldschmidt, S. On the Mechanism of Absorption from the Intestine. Physiol. Rev. 1921, 1 3(3), 421–453. 10.1152/physrev.1921.1.3.421.

(2) Magee, H. E. THE RÔLE OF THE SMALL INTESTINE IN NUTRITION. Physiol. Rev. 1930, 10 (3), 473–505. 10.1152/physrev.1930.10.3.473.

(3) Madara, J. L. Loosening Tight Junctions. Lessons from the Intestine. J. Clin. Invest. 1989, 83 (4), 1089–1094. 10.1172/JCI113987.

(4) Mobassaleh, M.; Montgomery, R. K.; Biller, J. A.; Grand, R. J. Development of Carbohydrate Absorption in the Fetus and Neonate. Pediatrics 1985, 75 (1), 160–166. 10.1542/peds.75.1.160.

(5) Moxey, P. C.; Trier, J. S. Specialized Cell Types in the Human Fetal Small Intestine. Anat. Rec. 1978, 191 (3), 269–285. 10.1002/ar.1091910302.

(6) Morson, B. C. Histopathology of the Small Intestine. Proc. R. Soc. Med. 1959, 52 (1), 6–10. 10.1177/003591575905200102.

(7) Wilson, J. P. Surface Area of the Small Intestine in Man. Gut 1967, 8 (6), 618–621. 10.1136/gut.8.6.618.

(8) Helander, H. F.; Fändriks, L. Surface Area of the Digestive Tract – Revisited. Scand. J. Gastroenterol. 2014, 49 (6), 681–689. 10.3109/00365521.2014.898326.

(9) Iqbal, J.; Hussain, M. M. Intestinal Lipid Absorption. Am J Physiol Endocrinol Metab 2009, 296 (6), E1183–E1194. 10.1152/ajpendo.90899.2008.

(10) Shirazi-Beechey, S. P. Molecular Biology of Intestinal Glucose Transport. Nutr. Res. Rev. 1995, 8 (1), 27–41. 10.1079/NRR19950005.

(11) Freeman, H. J.; Kim, Y. S. Digestion and Absorption of Protein. Annu. Rev. Med. 1978, 29 (1), 99–116. 10.1146/annurev.me.29.020178.000531.

(12) Bröer, S. Intestinal Amino Acid Transport and Metabolic Health. Annu. Rev. Nutr. 2023, 43 (1), 73–99. 10.1146/annurev-nutr-061121-094344.

(13) Rose, R. C. Water-Soluble Vitamin Absorption in Intestine. Annu. Rev. Physiol. 1980, 42 (1), 157–171. 10.1146/annurev.ph.42.030180.001105.

(14) Said, H.; Nexø, E. Gastrointestinal Handling of Water-Soluble Vitamins. In Comprehensive Physiology; 2018; Vol. 8, pp 1291–1311. 10.1002/cphy.c170054.

(15) Lin, L.; Yee, S. W.; Kim, R. B.; Giacomini, K. M. SLC Transporters as Therapeutic Targets: Emerging Opportunities. Nat Rev Drug Discov 2015, 14 (8), 543–560. 10.1038/nrd4626.

(16) George, A. M. ABC Transporters 45 Years On. Int J Mol Sci 2023, 24 (23), 16789. 10.3390/ijms242316789.

(17) Takayama, K.; Ito, K.; Matsui, A.; Yamashita, T.; Kawakami, K.; Hirayama, D.; Kishimoto, W.; Nakase, H.; Mizuguchi, H. In Vivo Gene Expression Profile of Human Intestinal Epithelial Cells: From the Viewpoint of Drug Metabolism and Pharmacokinetics. Drug Metab. Dispos. 2021, 49 (3), 221–232. 10.1124/dmd.120.000283.

(18) Drozdzik, M.; Busch, D.; Lapczuk, J.; Müller, J.; Ostrowski, M.; Kurzawski, M.; Oswald, S. Protein Abundance of Clinically Relevant Drug Transporters in the Human Liver and Intestine: A Comparative Analysis in Paired Tissue Specimens. Clin. Pharmacol. Ther. 2019, 105 (5), 1204–1212. 10.1002/cpt.1301.

(19) Mooij, M. G.; Schwarz, U. I.; Koning, B. A. E. de; Leeder, J. S.; Gaedigk, R.; Samsom, J. N.; Spaans, E.; Goudoever, J. B. van; Tibboel, D.; Kim, R. B.; Wildt, S. N. de. Ontogeny of Human Hepatic and Intestinal Transporter Gene Expression during Childhood: Age Matters. Drug Metab Dispos 2014, 42 (8), 1268–1274. 10.1124/dmd.114.056929.

(20) Liu, Y.; Beyer, A.; Aebersold, R. On the Dependency of Cellular Protein Levels on mRNA Abundance. Cell 2016, 165 (3), 535–550. 10.1016/j.cell.2016.03.014.

(21) Drozdzik, M.; Gröer, C.; Penski, J.; Lapczuk, J.; Ostrowski, M.; Lai, Y.; Prasad, B.; Unadkat, J. D.; Siegmund, W.; Oswald, S. Protein Abundance of Clinically Relevant Multidrug Transporters along the Entire Length of the Human Intestine. Mol. Pharm. 2014, 11 (10), 3547–3555. 10.1021/mp500330y.

(22) Couto, N.; Al-Majdoub, Z. M.; Gibson, S.; Davies, P. J.; Achour, B.; Harwood, M. D.; Carlson, G.; Barber, J.; Rostami-Hodjegan, A.; Warhurst, G. Quantitative Proteomics of Clinically Relevant Drug-Metabolizing Enzymes and Drug Transporters and Their Intercorrelations in the Human Small Intestine. Drug Metab Dispos 2020, 48 (4), 245–254. 10.1124/dmd.119.089656.

(23) Al-Majdoub, Z. M.; Couto, N.; Achour, B.; Harwood, M. D.; Carlson, G.; Warhurst, G.; Barber, J.; Rostami-Hodjegan, A. Quantification of Proteins Involved in Intestinal Epithelial Handling of Xenobiotics. Clin. Pharmacol. Ther. 2021, 109 (4), 1136–1146. 10.1002/cpt.2097.

(24) Kiss, M.; Mbasu, R.; Nicolaï, J.; Barnouin, K.; Kotian, A.; Mooij, M. G.; Kist, N.; Wijnen, R. M. H.; Ungell, A.-L.; Cutler, P.; Russel, F. G. M.; Wildt, S. N. de. Ontogeny of Small Intestinal Drug Transporters and Metabolizing Enzymes Based on Targeted Quantitative Proteomics. Drug Metab Dispos 2021, 49 (12), 1038–1046. 10.1124/dmd.121.000559.

(25) De Waal, T.; Handin, N.; Brouwers, J.; Miserez, M.; Hoffman, I.; Rayyan, M.; Artursson, P.; Augustijns, P. Expression of Intestinal Drug Transporter Proteins and Metabolic Enzymes in Neonatal and Pediatric Patients. Int. J. Pharm. 2024, 654, 123962. 10.1016/j.ijpharm.2024.123962.

(26) Goelen, J.; Farrell, G.; McGeehan, J.; Titman, C. M.; J. W. Rattray, N.; Johnson, T. N.; Horniblow, R. D.; Batchelor, H. K. Quantification of Drug Metabolising Enzymes and Transporter Proteins in the Paediatric Duodenum via LC-MS/MS Proteomics Using a QconCAT Technique. Eur. J. Pharm. Biopharm. 2023, 191, 68–77. 10.1016/j.ejpb.2023.08.011.

(27) Harder, F.; Brouwers, J.; Vanuytsel, T.; Oswald, S.; Augustijns, P. Advanced Age-Related Changes in Intestinal Drug Transporters and Metabolising Enzymes: A Targeted Proteomics Study. Mol. Pharm. 2026. 10.1021/acs.molpharmaceut.5c01755.

(28) Searle, B. C.; Pino, L. K.; Egertson, J. D.; Ting, Y. S.; Lawrence, R. T.; MacLean, B. X.; Villén, J.; MacCoss, M. J. Chromatogram Libraries Improve Peptide Detection and Quantification by Data Independent Acquisition Mass Spectrometry. Nat Commun 2018, 9 (1), 5128. 10.1038/s41467-018-07454-w.

(29) Govaert, E.; Van Steendam, K.; Willems, S.; Vossaert, L.; Dhaenens, M.; Deforce, D. Comparison of Fractionation Proteomics for Local SWATH Library Building. PROTEOMICS 2017, 17 (15–16), 1700052. 10.1002/pmic.201700052.

(30) Searle, B. C.; Swearingen, K. E.; Barnes, C. A.; Schmidt, T.; Gessulat, S.; Küster, B.; Wilhelm, M. Generating High Quality Libraries for DIA MS with Empirically Corrected Peptide Predictions. Nat Commun 2020, 11 (1), 1548. 10.1038/s41467-020-115346-1.

(31) Hughes, C. S.; Moggridge, S.; Müller, T.; Sorensen, P. H.; Morin, G. B.; Krijgsveld, J. Single-Pot, Solid-Phase-Enhanced Sample Preparation for Proteomics Experiments. Nat Protoc 2019, 14 (1), 68–85. 10.1038/s41596-018-0082-x.

(32) MacLean, B.; Tomazela, D. M.; Shulman, N.; Chambers, M.; Finney, G. L.; Frewen, B.; Kern, R.; Tabb, D. L.; Liebler, D. C.; MacCoss, M. J. Skyline: An Open Source Document Editor for Creating and Analyzing Targeted Proteomics Experiments. Bioinformatics 2010, 26 (7), 966–968. 10.1093/bioinformatics/btq054.

(33) Chambers, M. C.; Maclean, B.; Burke, R.; Amodei, D.; Ruderman, D. L.; Neumann, S.; Gatto, L.; Fischer, B.; Pratt, B.; Egertson, J.; Hoff, K.; Kessner, D.; Tasman, N.; Shulman, N.; Frewen, B.; Baker, T. A.; Brusniak, M.-Y.; Paulse, C.; Creasy, D.; Flashner, L.; Kani, K.; Moulding, C.; Seymour, S. L.; Nuwaysir, L. M.; Lefebvre, B.; Kuhlmann, F.; Roark, J.; Rainer, P.; Detlev, S.; Hemenway, T.; Huhmer, A.; Langridge, J.; Connolly, B.; Chadick, T.; Holly, K.; Eckels, J.; Deutsch, E. W.; Moritz, R. L.; Katz, J. E.; Agus, D. B.; MacCoss, M.; Tabb, D. L.; Mallick, P. A Cross-Platform Toolkit for Mass Spectrometry and Proteomics. Nat Biotechnol 2012, 30 (10), 918–920. 10.1038/nbt.2377.

(34) Gessulat, S.; Schmidt, T.; Zolg, D. P.; Samaras, P.; Schnatbaum, K.; Zerweck, J.; Knaute, T.; Rechenberger, J.; Delanghe, B.; Huhmer, A.; Reimer, U.; Ehrlich, H.-C.; Aiche, S.; Kuster, B.; Wilhelm, M. Prosit: Proteome-Wide Prediction of Peptide Tandem Mass Spectra by Deep Learning. Nat Methods 2019, 16 (6), 509–518. 10.1038/s41592-019-0426-7.

(35) Lautenbacher, L.; Yang, K. L.; Kockmann, T.; Panse, C.; Chambers, M.; Kahl, E.; Yu, F.; Gabriel, W.; Bold, D.; Schmidt, T.; Li, K.; MacLean, B.; Nesvizhskii, A. I.; Wilhelm, M. Koina: Democratizing Machine Learning for Proteomics Research. bioRxiv 2024, 2024.06.01.596953. 10.1101/2024.06.01.596953.

(36) Käll, L.; Canterbury, J. D.; Weston, J.; Noble, W. S.; MacCoss, M. J. Semi-Supervised Learning for Peptide Identification from Shotgun Proteomics Datasets. Nat Methods 2007, 4 (11), 923–925. 10.1038/nmeth1113.

(37) Choi, M.; Chang, C.-Y.; Clough, T.; Broudy, D.; Killeen, T.; MacLean, B.; Vitek, O. MSstats: An R Package for Statistical Analysis of Quantitative Mass Spectrometry-Based Proteomic Experiments. Bioinformatics 2014, 30 (17), 2524–2526. 10.1093/bioinformatics/btu305.

(38) Sharma, V.; Eckels, J.; Schilling, B.; Ludwig, C.; Jaffe, J. D.; MacCoss, M. J.; MacLean, B. Panorama Public: A Public Repository for Quantitative Data Sets Processed in Skyline. Mol Cell Proteomics 2018, 17 (6), 1239–1244. 10.1074/mcp.RA117.000543.

(39) Vizcaíno, J. A.; Deutsch, E. W.; Wang, R.; Csordas, A.; Reisinger, F.; Ríos, D.; Dianes, J. A.; Sun, Z.; Farrah, T.; Bandeira, N.; Binz, P.-A.; Xenarios, I.; Eisenacher, M.; Mayer, G.; Gatto, L.; Campos, A.; Chalkley, R. J.; Kraus, H.-J.; Albar, J. P.; Martinez-Bartolomé, S.; Apweiler, R.; Omenn, G. S.; Martens, L.; Jones, A. R.; Hermjakob, H. ProteomeXchange Provides Globally Coordinated Proteomics Data Submission and Dissemination. Nat Biotechnol 2014, 32 (3), 223–226. 10.1038/nbt.2839.

(40) Becker, H. M.; Seidler, U. E. Bicarbonate Secretion and Acid/Base Sensing by the Intestine. Pflugers Arch - Eur J Physiol 2024, 476 (4), 593–610. 10.1007/s00424-024-02914-3.

(41) Murata, M.; Fujioka, H.; Yokota, J.; Okada, K.; Yamashita, T.; Hirayama, D.; Kawakami, K.; Morinaga, G.; Saito, A.; Nakase, H.; Kishimoto, W.; Mizuguchi, H. Regional Transcriptomics and Proteomics of Pharmacokinetics-Related Genes in Human Intestine. Mol. Pharm. 2023, 20 (6), 2876–2890. 10.1021/acs.molpharmaceut.2c01002.

(42) Zhang, H.; Wolford, C.; Basit, A.; Li, A. P.; Fan, P. W.; Murray, B. P.; Takahashi, R. H.; Khojasteh, S. C.; Smith, B. J.; Thummel, K. E.; Prasad, B. Regional Proteomic Quantification of Clinically Relevant Non-Cytochrome P450 Enzymes along the Human Small Intestine. Drug Metab. Dispos. 2020, 48 (7), 528–536. 10.1124/dmd.120.090738.

(43) Varum, F. J. O.; Veiga, F.; Sousa, J. S.; Basit, A. W. Mucus Thickness in the Gastrointestinal Tract of Laboratory Animals. J Pharm Pharmacol 2012, 64 (2), 218–227. 10.1111/j.2042-7158.2011.01399.x.

(44) Mouly, S.; Paine, M. F. P-Glycoprotein Increases from Proximal to Distal Regions of Human Small Intestine. Pharm Res 2003, 20 (10), 1595–1599. 10.1023/A:1026183200740.

(45) Scalise, M.; Galluccio, M.; Console, L.; Pochini, L.; Indiveri, C. The Human SLC7A5 (LAT1): The Intriguing Histidine/Large Neutral Amino Acid Transporter and Its Relevance to Human Health. Front. Chem. 2018, 6. 10.3389/fchem.2018.00243.

(46) Meier, C.; Camargo, S. M.; Hunziker, S.; Moehrlen, U.; Gros, S. J.; Bode, P.; Leu, S.; Meuli, M.; Holland-Cunz, S.; Verrey, F.; Vuille-dit-Bille, R. N. Intestinal IMINO Transporter SIT1 Is Not Expressed in Human Newborns. Am. J. Physiol. Gastrointest. Liver. Physiol. 2018, 315 (5), G887–G895. 10.1152/ajpgi.00318.2017.

(47) Malo, C. Multiple Pathways for Amino Acid Transport in Brush Border Membrane Vesicles Isolated from the Human Fetal Small Intestine. Gastroenterology 1991, 100 (6), 1644–1652. 10.1016/0016-5085(91)90664-7.

