## Supplemental Material for "Proteomic profiling of drug and nutrient transporter expression by small intestinal region in neonates, pediatrics, and adults"

### Methods

#### Small intestine dissection

Duodenum was identified by the presence of adjacent pancreatic tissue (if present), circular folds, a wider diameter relative to the jejunum or ileum, and the duodenojejunal flexure (if present). Jejunum was identified by a smaller diameter relative to the duodenum and the presence of circular folds that were at higher density than in the duodenum. Ileum was identified by the presence of a smaller diameter relative to the jejunum, sparse circular folds, the ileocecal junction (if present), and the presence of adjacent cecum/appendix tissue (if present).

Donor exclusion criteria included hepatitis B (antigen, core, NAT) positive serology, hepatitis C positive serology, HIV positive serology, bacterial or viral meningitis, MRSA, RPR/syphilis positive serology, sepsis, and approximate cold ischemic time > 24 hours. Exclusion criteria were implemented prior to sample collection.

#### Kingfisher sample preparation protocol

**Reagent info**

| Plate | Contents | Well volume (µL) |
|---|---|---|
| 1-Protein Capture | 100% Acetonitrile | 520 |
| | Lysate | 210.3 |
| | Beads | 12.5 |
| 2-ACN Wash 1 | 95% Acetonitrile | 1000 |
| 3-ACN Wash 2 | 95% Acetonitrile | 1000 |
| 4-ACN Wash 3 | 95% Acetonitrile | 1000 |
| 5-EtOH Wash 1 | 70% Ethanol | 1000 |
| 6-EtOH Wash 2 | 70% Ethanol | 1000 |
| 7-Trypsin Elution | 50 mM ABC, Trypsin 1:20 | 50 |
| Tip Plate | n/a | n/a |

**Steps Data**

| Step | Plate | Details |
|---|---|---|
| Pick-Up | Tip Plate | n/a |
| Bind Proteins | P1-Protein Capture | |
| Beginning of step | Precollect | No |
| Mixing/heating | Shake 1 time, speed | 00:01:00, Medium |
| | Shake 2 time, speed | 00:10:00, Paused |
| | Loop Count | 2 |
| | Tip position when paused | Tip edge in liquid |
| | Heating during mixing | No |
| End of step | Postmix | No |
| | Collect count | 5 |

| | | |
|---|---|---|
| | Collect time (s) | 30 |
| Wash 1 | P2-ACN Wash 1 | |
| Beginning of step | Precollect | No |
| | Release beads | No |
| Mixing/heating | Mixing time, speed | 00:02:30, Slow |
| | Heating during mixing | No |
| End of step | Postmix | No |
| | Collect beads | No |
| Wash 2 | P3-ACN Wash 2 | |
| Beginning of step | Precollect | No |
| | Release beads | No |
| Mixing/heating | Mixing time, speed | 00:02:30, Slow |
| | Heating during mixing | No |
| End of step | Postmix | No |
| | Collect beads | No |
| Wash 3 | P4-ACN Wash 3 | |
| Beginning of step | Precollect | No |
| | Release beads | No |
| Mixing/heating | Mixing time, speed | 00:02:30, Slow |
| | Heating during mixing | No |
| End of step | Postmix | No |
| | Collect beads | No |
| Wash 4 | P5-EtOH Wash 1 | |
| Beginning of step | Precollect | No |
| | Release beads | No |
| Mixing/heating | Mixing time, speed | 00:02:30, Slow |
| | Heating during mixing | No |
| End of step | Postmix | No |
| | Collect beads | No |
| Wash 5 | P6-EtOH Wash 2 | |
| Beginning of step | Precollect | No |
| | Release beads | No |
| Mixing/heating | Mixing time, speed | 00:02:30, Slow |
| | Heating during mixing | No |
| End of step | Postmix | No |
| | Collect beads | No |
| Pause | P7-Trypsin Elution | |
| | Message | Add digestion/elution plate |
| Protein Digestion and Elution | P7-Trypsin Elution | |
| Beginning of step | Precollect | No |
| | Release time, speed | 00:00:20, Bottom mix |
| Mixing/heating | Shake 1 time, speed | 00:00:15, Medium |
| | Shake 2 time, speed | 00:02:15, Paused |
| | Loop Count | 24 |
| | Tip position when paused | Tip edge in liquid |

| | | |
|---|---|---|
| | Heating temperature (˚C) | 37 |
| | Preheat | Yes |
| End of step | Postmix | No |
| | Collect count | 5 |
| | Collect time (s) | 30 |
| Dispose Beads | P1-Protein Capture | |
| | Release time, speed | 00:00:30, Medium |
| Leave | Tip Plate | |

**Matched-matrix calibration curve**

Yeast (Saccharomyces cerevisiae, Type II (Baker's yeast), Sigma Aldrich, St. Louis, MO) was lysed by sonication (Fisher Scientific sonic dismembrator model 100, set to continuous at 4 or 5, 10-second intervals for 1 minute) in 500 µL 20 mM Tris, 50 mM mannitol, 1 mM EDTA buffer at pH 7.4 (adjusted with HCl) with protease inhibitor. Protein concentration in yeast lysate was estimated with the BCA assay and aliquots of yeast lysate were digested as described in the Methods (Sample preparation and Protein digestion sections).

Standards were prepared by diluting digested, pooled mucosa samples (after sample reconstitution, described in Methods under Peptide sample analysis by DIA-MS) in digested yeast lysate. The following standards (expressed as % of pure pooled mucosa) were prepared in each calibration curve: 0.5, 1, 5, 10, 25, 50, 70, and 100%. Each standard had a total peptide concentration of 0.5 µg/µL in the reconstitution solution below. Calibration curves were prepared in technical triplicate and each curve was injected from low to high mucosa percentage once on the mass spectrometer.

**Chromatogram library generation**

Equal volumes of digested mucosa samples and 1 µL of digested yeast enolase were combined to generate a chromatogram library sample. A chromatogram library was generated by gas-phase fractionation of 1 µg of library sample on-column with narrow-window DIA runs of 4 m/z windows (with overlapping by 2 m/z)[30]. Each gas-phase fraction spanned 100 m/z with a total of 6 fractions spanning 395-1005 m/z. Narrow-window DIA runs were acquired (25 second expected precursor peak width and +3 charge, ion accumulation time of 23 ms, positive polarity, NCE of 27, 30,000 resolution, automatic gain control target of 1,000,000) with two MS scans to acquire precursor spectra every 25 MS/MS spectra. One MS scan covered the target m/z range ± 5 m/z (e.g., 395-505 m/z) while the other MS scan spanned 400-1600 m/z. Both MS scans had an expected precursor peak width of 25 seconds and charge of 3, ion accumulation time of 23 ms, positive polarity, 60,000 resolution, and an automatic gain control target of 1,000,000.

**Quantitative peptide identification**

Quantitative peptides were identified for all small intestine marker proteins, SLC/ABC transporters highlighted in dynamic range plots, and SLC/ABC transporters that were differentially expressed (see Results). Quantitative peptides for these select proteins were identified by examining a bi-linear regression fit to the matched-matrix calibration curves in Skyline.

### Supplemental Figures and Tables

### Table S1. Donor demographics and sample information.

| Sample Identifier | Sex | Age | Race | Cause of death | Notes |
|---|---|---|---|---|---|
| HINT 2 | F | 5 days old | White (Hispanic) | Anoxia, Cardiovascular (maternal uterine rupture) | n/a |
| HID04 | M | 1 month old | White | Anoxia, Cardiovascular (shoulder dystocia) | Only jejunum received |
| PHI4 | M | 1 month old | Black or African American | Anoxia, Cardiovascular (aspiration) | Only duodenum and jejunum received |
| PHI5 | M | 12 months old | White | Head Trauma (MVA) | n/a |
| HINT 3 | M | 2 years old | White | Unknown (natural causes) | n/a |
| PHI2 | F | 2 years old | not reported | Anoxia, Cardiovascular (non-MVA) | Only ileum analyzed |
| PHI3 | M | 4 years old | White | Head Trauma (MVA) | n/a |
| HINT 1 | M | 5 years old | African American | Drug-induced toxicity | n/a |
| PHI6 | M | 5 years old | White | Anoxia (drowning) | n/a |
| PHI1 | F | 8 years old | Black or African American | Anoxia, Cardiovascular (cardiac) | Only jejunum and ileum received |
| AHI7 | M | 18 years old | White | Anoxia (marijuana/opioid intoxication) | n/a |
| AHI2 | M | 20 years old | White | Anoxia, Cardiovascular (natural causes) | Only jejunum received |
| AHI8 | F | 23 years old | White | Anoxia (tylenol intoxication) | n/a |
| AHI4 | F | 24 years old | White | Cerebrovascular/stroke (intracranial hemorrhage/stroke) | Only ileum received |
| AHI6 | F | 27 years old | White | Anoxia, Cardiovascular | n/a |
| AHI5 | F | 32 years old | White | CNS tumor | Only jejunum and ileum received |
| AHI1 | M | 34 years old | African American | Anoxia, Cardiovascular (natural causes) | Only jejunum and ileum received; donor had diabetes |
| AHI3 | M | 34 years old | White | Anoxia, Cardiovascular (natural causes) | Only jejunum and ileum received; donor had pancreatitis |

F: female, M: male, MVA: motor-vehicle accident, n/a: not applicable

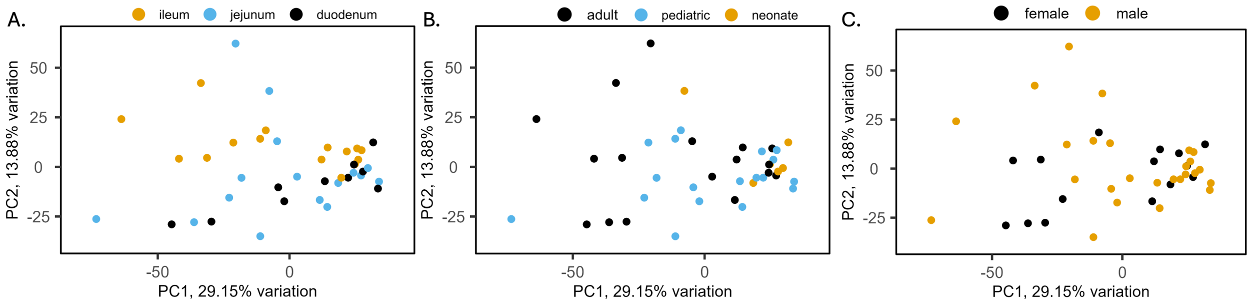


**Figure S1. PCA of samples by intestinal region, age, or sex.** Samples cluster by proximal small intestine regions (duodenum or jejunum) or ileum (A) but not strongly by age group (B). No clustering by donor sex was observed (C). PC = principal component

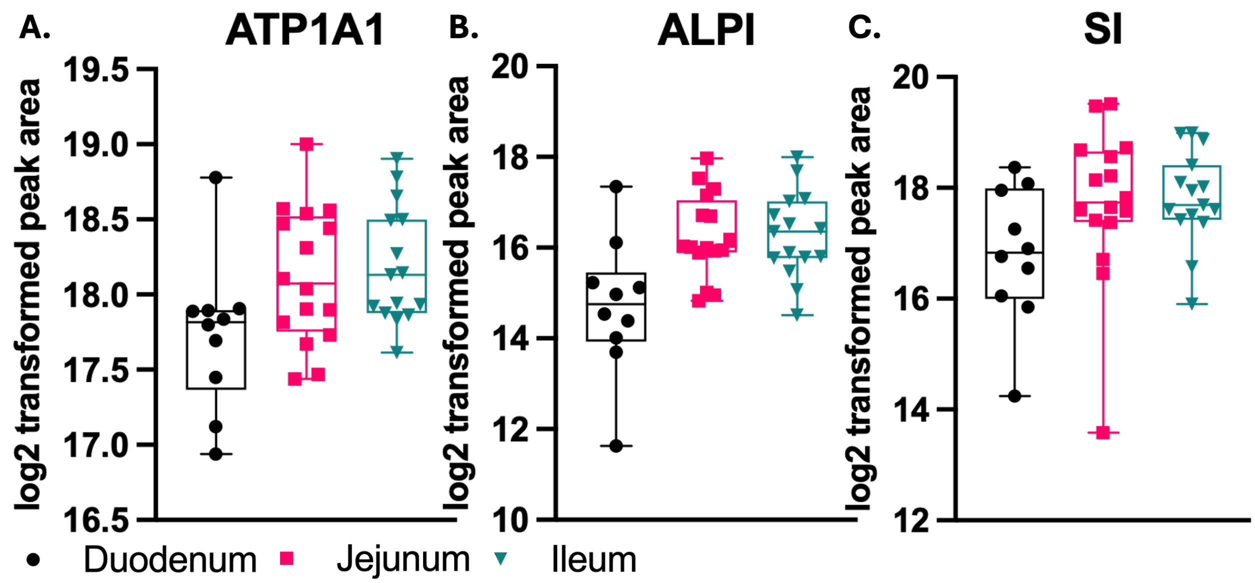


**Figure S2. Small intestine marker proteins by intestinal region.** The small intestine marker proteins ATP1A1 (A), ALPI (B), and SI (C) did not show trends with small intestine region.

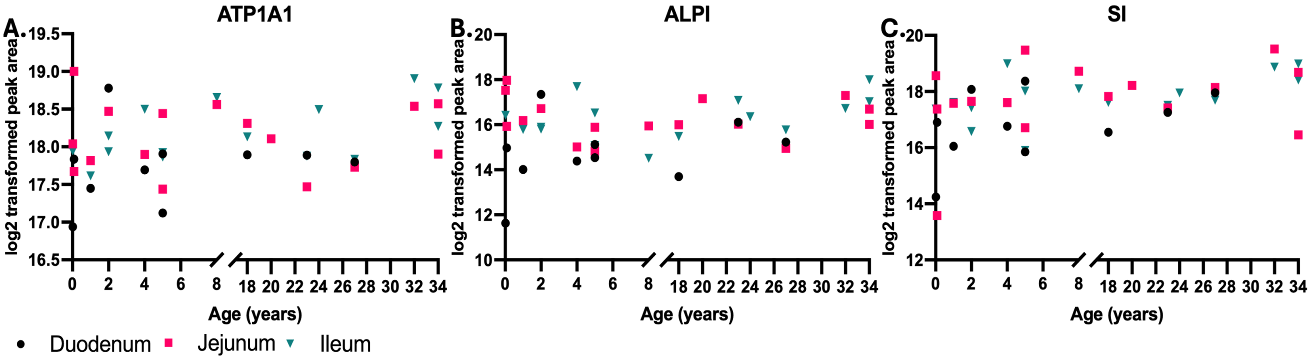


**Figure S3. Small intestine marker proteins by age.** ATP1A1 (A), ALPI (B), and SI (C) did not have a significant trends by small intestinal region with age.

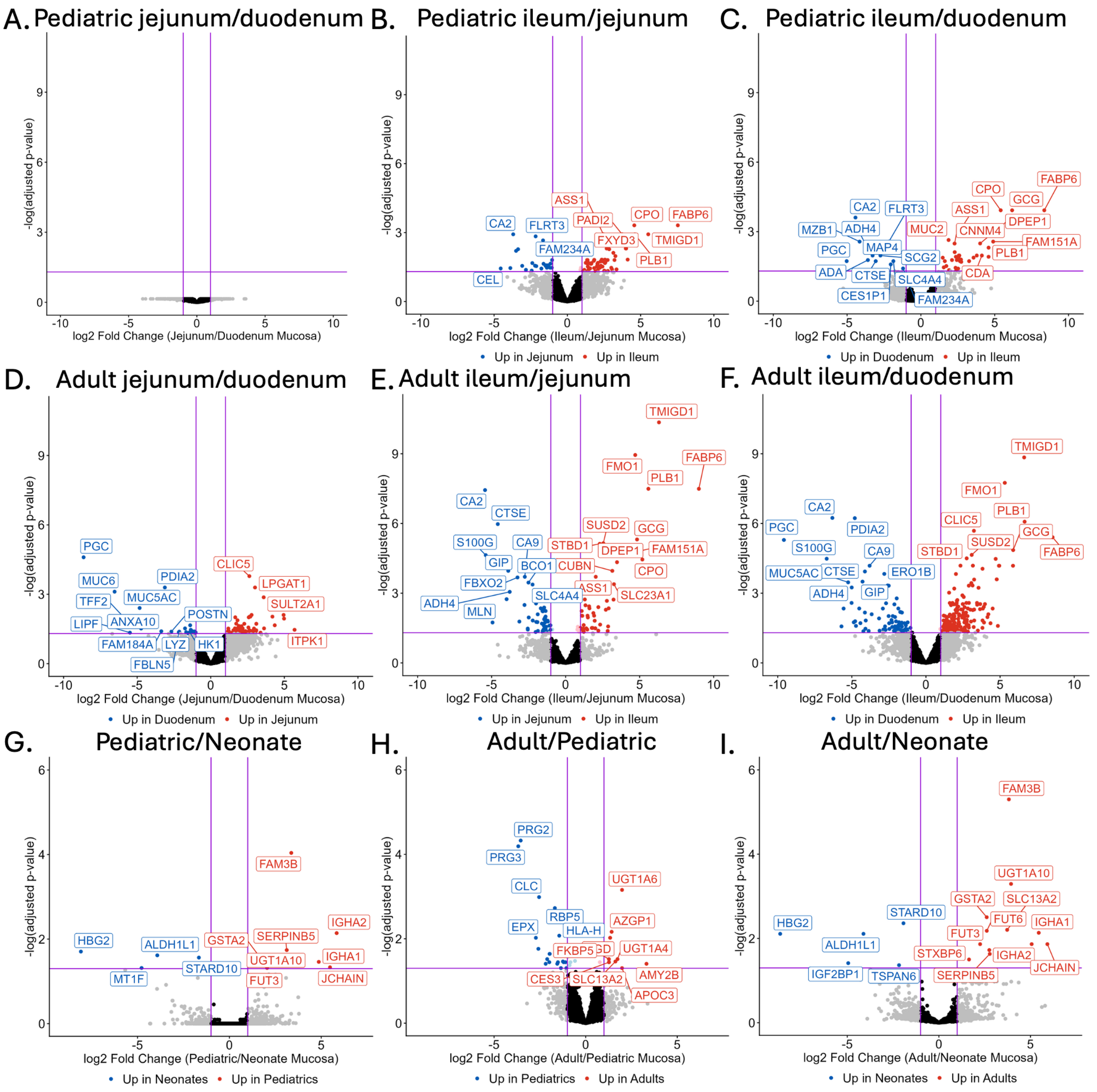


**Figure S4. Differentially expressed proteins by intestinal region or age.** Pediatric regions (A-C), adult regions (D-F), age groups (G-I). No SLC or ABC transporters are differentially expressed between neonates and pediatrics (G), but SLC13A2 is significantly higher with age in adults compared to pediatrics (H) or neonates (I). n = 5 pediatric duodenum, n = 6 pediatric jejunum, n = 7 pediatric ileum, n = 3 adult duodenum, n = 7 adult jejunum, n = 7 adult ileum; t-tests with multiple testing corrected by Benjamini-Hochberg method; pediatric duodenum versus jejunum n = 5,762 proteins, pediatric jejunum versus ileum n = 5,784 proteins, pediatric duodenum versus ileum n = 5,759 proteins, adult duodenum versus jejunum n = 5,657 proteins, pediatric jejunum versus ileum n = 5,771 proteins, pediatric duodenum versus ileum n = 5,648 proteins, pediatric versus neonate n = 5,725 proteins, adult versus pediatric n = 5,867 proteins, adult versus neonate n = 5,725 proteins
